# Sustained LDL-C Lowering Reveals Selective Expansion of Fibroblast Subpopulations with Distinct Regulatory and Matrix Programs During Atherosclerosis Remodeling

**DOI:** 10.64898/2026.11.25.768950

**Authors:** Johana Coronel, Alexander C. Bashore, Chenyi Xue, Guanliang Li, Enrique J. García, Lucie Y. Zhu, Shareef Khalid, Wenli Liu, Brian D. Hardaway, Ira Tabas, Alan R. Tall, Judith C. Sluimer, Muredach P. Reilly

**Author notes:** Corresponding author: Muredach P. Reilly.

## Abstract

**Background:** Smooth muscle cells (SMCs) and SMC-derived cells contribute to atherosclerotic plaque stability through matrix production and fibrous cap formation. Recent studies have also identified distinct fibroblast populations in atherosclerotic lesions, but how these populations respond to sustained LDL-C lowering and contribute to plaque remodeling remains poorly understood.

**Methods:** We combined serial histological assessment, SMC lineage tracing, and single-cell RNA sequencing to examine vascular stromal cell and lesion structural responses during 20 weeks of sustained LDL-C lowering in mice with established atherosclerosis. Regulatory network analysis and computational perturbation predictions were supported by KLF3 and FOXD1 silencing in cultured mouse and human aortic fibroblasts.

**Results:** LDL-C lowering led to rapid reductions in macrophage content and cleaved gasdermin D, followed by decreased necrotic core area and increased collagen content and fibrous cap thickness, with minimal change in lesion area. During this remodeling, SMC-derived cell populations changed little. In striking contrast, CD26⁺ and FMOD⁺ fibroblast subtypes increased by 5 weeks and remained elevated through 20 weeks, with increased fibroblast-associated signals in the intima and fibrous cap. These populations showed distinct matrix programs, with ECM glycoprotein and regulatory programs enriched in CD26⁺ fibroblasts and collagen and proteoglycan programs in FMOD⁺ fibroblasts. Regulatory network analysis identified KLF3 and FOXD1 as candidate master regulators of CD26⁺ and FMOD⁺ fibroblasts, respectively. Silencing either factor reduced selected matrix-associated and proliferation transcripts and impaired scratch closure in mouse and human aortic fibroblasts, while KLF3 silencing also increased selected inflammatory genes. Exploratory human genetic analyses identified associations of KLF3-region and FOXD1 variation with coronary artery disease.

**Conclusions:** These findings reveal fibroblast subtype-specific responses during sustained LDL-C lowering and identify KLF3 and FOXD1 as candidate transcriptional regulators of fibroblast matrix-associated and inflammatory functions relevant to plaque remodeling and stabilization.

## Introduction

Atherosclerotic cardiovascular disease (CVD) remains a leading cause of morbidity and mortality worldwide^1^. Although statins and PCSK9 inhibitors substantially lower low-density lipoprotein cholesterol (LDL-C) and reduce cardiovascular events, considerable residual CVD risk persists^2, 3^. Plaque rupture contributes to acute cardiovascular events, highlighting the importance of understanding the cellular mechanisms that promote plaque stability and prevent rupture^4, 5^. Stable plaques are associated with thick, collagen-rich fibrous caps, smaller necrotic cores, and less inflammation than rupture-prone plaques^4^. Intensive LDL-C lowering promotes these clinically relevant features of plaque stability^6, 7^. However, the cellular and molecular processes responsible for this remodeling remain incompletely understood.

Smooth muscle cells (SMCs) and SMC-derived cells contribute to plaque stability by producing extracellular matrix (ECM), including collagen within the fibrous cap^8, 9^. Lineage-tracing and single-cell studies have shown that SMC phenotypic modulation generates matrix-producing cells with fibroblast-like characteristics^8, 10^. Yet, these SMC-derived fibroblast cells do not account for all plaque fibroblasts like cells-we and others have shown recently that authentic fibroblast populations of non SMC-lineage are also present in lesions^11^. Our group identified a CD26⁺ fibroblast subset in the adventitia and lesion intima during atherosclerosis progression^12^. Other studies have defined vascular fibroblast populations with different matrix-associated and inflammatory programs^11, 13, 14^. Lineage tracing in mouse models showed that most plaque fibroblasts were *Myh11*-lineage negative and that *Pdgfra*-lineage-positive cells largely retained fibroblast identity during disease progression^11^. Together, these findings establish fibroblasts as a distinct and heterogeneous component of the plaque stromal compartment.

Much of this recent evidence comes from experimental mouse studies of atherosclerosis progression or cross-sectional interrogation of human plaques^10–14^. Studies of disease progression and cross-sectional human plaques establish fibroblast heterogeneity but do not determine how their abundance and function change during LDL-cholesterol lowering induced lesion regression^11–14^. Fibroblasts produce and remodel ECM during tissue responses to injury and can also regulate local inflammation^7, 15^, providing a biological rationale for their contribution to plaque stabilization during LDL-C lowering. Thus, key knowledge gaps are how fibroblast populations and their matrix programs change during sustained lipid lowering and which transcriptional regulators control these programs. Fibroblast matrix-associated and inflammatory functions also raise the possibility that fibroblast–immune communication contributes to lesion remodeling.

In the present study, we combined serial histological assessment, single-cell RNA sequencing (scRNA-seq) and SMC lineage tracing, in the first longitudinal analysis of prolonged vascular stromal cell dynamics-both SMC-derived and fibroblast-and lesion structural responses during 20 weeks of lipid lowering in mice with established atherosclerosis. This is the first approach enabled us to capture dynamic changes in cell compositions during both the early inflammatory resolution and the subsequent structural remodeling associated with plaque stabilization and healing. LDL-C lowering led to a rapid reduction in inflammatory burden. We distinguished fibroblasts from SMC-derived cells and identified major increases in CD26⁺ and FMOD⁺ fibroblasts with different matrix-associated gene programs while SMC-derived cells did not change. Transcription factor activity estimates and computational perturbation predictions identified KLF3 and FOXD1 as candidate master regulators of CD26⁺ and FMOD⁺ fibroblasts respectively. Depletion studies in cultured mouse and human aortic fibroblasts demonstrated distinct effects of KLF3 and FOXD1 depletion on matrix-associated gene expression, while KLF3 knockdown also increased selected inflammatory mediators. These findings reveal novel fibroblast subtype-specific responses in lesions during LDL-C lowering and identify KLF3 and FOXD1 as candidate transcriptional regulators of fibroblast matrix-associated and inflammatory functions in plaque remodeling and stabilization after LDL-C lowering.

## Methods

### Mouse Studies

All animal experiments were conducted in compliance with the Institutional Animal Care and Use Committee (IACUC) of Columbia University under protocol number AABQ5576. Mouse strains were obtained from The Jackson Laboratory, including Ldlr KO (Stock: 002207), ZsGreen (Stock: 007906), and Myh11-CreERT2 (Stock: 019079). These strains were bred on a C57BL/6J background to create a hyperlipidemic smooth muscle cell lineage tracing model: Ldlr KO; ZsGreen; Myh11-CreER^T^^210, 16, 17^. Only male mice were used due to the insertion of the Myh11-CreERT2 transgene on the Y chromosome. To activate Cre recombinase for ZsGreen lineage tracing, mice received tamoxifen (40 mg/kg body weight, intraperitoneally) once daily for 5 consecutive days to induce permanent ZsGree1 labeling of Myh11-lineage cells. A washout period of at least 10 days was observed after tamoxifen administration. Atherosclerosis was induced by feeding mice a Western Diet (WD) containing 40% Kcal of fat and 0.15% Cholesterol (Research Diet, D12079Bi) for 16 weeks. A subset of mice was sacrificed at the end of WD feeding to assess baseline atherosclerosis. The remaining mice underwent atherosclerosis regression by switching to a chow diet for 2, 5, 10, or 20 weeks and received a helper-dependent adenovirus encoding human *Ldlr* gene (HDAd-LDLR) by tail injection. This non-integrating viral vector targets the liver, restoring *Ldlr* expression in hepatocytes and reducing circulating cholesterol in *Ldlr^-/-^*mice^18, 19^.

### Blood Sampling and Plasma Cholesterol Measurement

Mice were anesthetized with isoflurane, and blood was collected by retroorbital puncture into heparinized microcapillary tubes. Blood samples were collected before HDad-LDLR administration and during regression, including 2 days, and 1, 2, 3, 4, 5, 10, 15, and 20 weeks after HDAd-LDLR injection to monitor LDL-C reductions. Plasma was separated by centrifugation at 1500 x g for 10 minutes at 4C and stored at -80C. Total plasma cholesterol was measured using Infinity Cholesterol Reagent (Thermo Fisher Scientific, TR13421) according to the manufacturer’s protocol.

### Tissue Collection, Fixation, sectioning, and Lesion Analysis

At the experimental endpoint, mice were euthanized and perfused with phosphate-buffered saline (PBS). Hearts containing the aortic root were collected, fixed in 4% paraformaldehyde for 24 hours, transferred to 70% ethanol, and embedded in paraffin. Blocks were sectioned at 6 µm using a Leica RM2235 microtome. Sections were stained with hematoxylin and eosin to quantify lesion and necrotic core areas and with Picrosirius Red to assess collagen content and fibrous-cap thickness. CD68 immunostaining was used to quantify macrophage content, and cleaved gasdermin D immunofluorescence was quantified as mean fluorescence intensity within the lesion. Images were analyzed in ImageJ/Fiji by an investigator blinded to experimental group. Collagen-positive and CD68-positive areas were expressed as percentages of total lesion area, whereas cleaved gasdemin D fluorescence was expressed as mean fluorescence intensity within the lesion area. For each endpoint, multiple aortic root sections per mouse were analyzed, and measurements were averaged to obtain a single value per mouse for statistical analysis. Hematoxylin and eosin and Picrosirius Red staining were performed by the Columbia University Molecular Pathology Shared Resource.

### Aortic Digestion for Single Cell-RNA Sequencing (scRNA-seq)

At sacrifice, the vasculature was perfused with ice-cold DPBS containing 1 µg/mL Actinomycin D (Thermo Fisher Scientific, 11805017). The aorta, including the ascending aorta, brachiocephalic artery (BCA), and thoracic aorta, were harvested and digested in RPMI supplemented with 4 U/mL Liberase (Roche, 05401119001), 60 U/mL hyaluronidase (Millipore Sigma, H3506-100MG), and 60 U/mL DNase (Worthington, LS006331). The resulting cell suspension was washed with FACS buffer containing 2% HI-FBS, 5 mM EDTA, 20 mM HEPES, 1 mM Sodium Pyruvate in 1X DPBS and centrifuged at 400 x g for 5 minutes at 4C. Cells were then resuspended in 49 µL of FACS buffer, and 1 µL of TruStain FcX PLUS (anti-mouse CD16/32) (BioLegend, 156603) was added for 10 minutes at 4C to block Fc receptors. The TotalSeq-A Mouse Universal Cocktail (BioLegend, 199901) oligo-conjugated antibodies were reconstituted in 100 µL of FACS buffer, and 30 µL of this cocktail was added to each single-cell suspension, incubating for 30 minutes at 4C. Samples were washed twice in FACS buffer with centrifugation at 400g for 5 minutes at 4C. For cell hashing, cell pellets were resuspended in Cell Multiplexing Oligos (10x Genomics) and incubated for 5 minutes at room temperature, followed by three washes in FACS buffer with centrifugation at 400g for 5 minutes at 4C. Single-cell suspensions were stained with DAPI (1:3,000) and DRAQ5 (1:1,000). Viable cells, defined as DRAQ5-positive and DAPI-negative, were sorted using a BD FACSAria II and collected into 1.5 mL Eppendorf tubes containing DMEM/F12 + 10% FBS. Cells were then pelleted and counted for input into the 10x Genomics Chromium system.

### ScRNA-seq Data Processing and Analyses

Fastq files were processed by “cell ranger multi” pipeline from Cell Ranger 6.1.2. Each sequencing run contained pooled libraries from 3 to 4 replicates, each tagged with multiplexing oligos that were specified in the pipeline configuration file. Reads were mapped to a modified mouse genome with ZsGreen1 sequence^17^ appended to genome build GRCm38. Transcriptome mapping was based on GENCODE vM23 annotation with the addition of ZsGreen1 annotation. Count matrices of unique molecular identifier (UMI) were generated for each replicate separately. A suffix number was added to gene symbols when multiple symbols share the same gene ID.

Gene and cell level filtering were performed on each replicate using protocols we previously published^17^. Briefly, as replicates from each sequencing run have consistent distributions of UMI counts, gene counts, and mitochondrial content, the same filtering parameters were applied to replicates in a sequencing run. Genes were included if they were present in ≥10 cells. Cells were retained if they had ≥200 genes expressed. A maximum gene count filter and a maximum UMI count filter were applied to cells. The parameters varied slightly due to different sequencing output and are summarized in **Supp. Table A1**. A maximum of 10% reads mapped to mitochondrial genes were allowed in each cell. Filtered UMI count matrices from all-time points were integrated and clustered using CarDEC^20^. Run ID was used as the batch variable, and 2,000 highly variable genes were selected for integration and clustering. A total of 21 clusters were identified in CarDEC. Differential expression analysis was conducted for each cluster compared to other cells using Wilcoxon rank-sum test in Scanpy^21^. UMI counts were first normalized by total counts across genes with the target sum set to 10,000 and then transformed by natural logarithm. Genes with fold change ≥1.5, FDR adjusted P-value <0.05, and presence in ≥25% of cells in either group were considered differentially expressed. Clusters were annotated manually using top features from each cluster and known cell type markers in atherosclerosis context. Four smooth muscle cell (SMC) populations were collapsed into one SMC population as they had high *Myh11* expression and shared similar top differentially expressed genes. A modulated SMC population with 62.4% of cells being ZsGreen1-negative were split into two populations by ZsGreen1 status: chondromyocyte, characterized by its SMC origin and expression of chondrocyte genes such as *Col2a1* and *Spp1,* and myofibroblast, characterized by its non-SMC origin and expression of contractile SMC genes such as Myh11 and Acta2 as well as fibroblast genes such as *Dcn*. These subpopulations names are used consistently throughout the manuscript.

Macrophages identified in the main analysis were extracted for deeper sub clustering analysis using CarDEC. 2,000 highly variable genes were selected and run ID was specified as the batch variable. A total of 9 clusters were identified, including 5 macrophage subpopulations, 2 dendritic cell populations, a small contaminating fibroblast population, and a monocyte population.

### Over-representation Analysis and Gene set Enrichment Analysis

ClusterProfiler^22^ was used for over-representation analysis (ORA) of differentially expressed genes in each of the following gene sets: Gene Ontology (GO) biological pathway, KEGG pathway (release 117.0), and Reactome pathway. In addition, mouse matrisome gene sets characterized by Naba et al.^23^ were downloaded from mouse MSigDB^24, 25^ and tested for over-representation. Gene symbols were converted to Entrez IDs before analysis. P-values were adjusted by Benjamini-Hochberg^26^ approach. Redundant over-represented gene sets from GO terms were simplified using the “simplify” function in clusterProfiler with similarity cutoff set to 0.7. In scRNA-seq, ORA was performed separately for genes that are upregulated in FMOD fibroblasts and upregulated in CD26 fibroblasts. Background genes were restricted to genes expressed in either of the two populations. Gene set enrichment analysis (GSEA) was performed using the same gene sets as in ORA. In scRNA-seq, 5,094 genes were included that were expressed in ≥10% of cells in CD26⁺ or FMOD⁺ fibroblasts and were sorted by log2 fold change. P-values were adjusted by Benjamini-Hochberg approach.

### Master Regulator Analysis

Master regulator analysis was applied to scRNA-seq fibroblast populations to identify regulators that were activated in one fibroblast population compared to the others. Data processing, network generation, and protein activity inference followed the guidelines in PISCES pipeline^27^. UMI counts were extracted from CD26⁺ fibroblast, FMOD⁺ fibroblast, and myofibroblast populations. MetaCells were generated by first randomly selecting 250 cells from each of the CD26⁺ fibroblast and FMOD⁺ fibroblast populations and then aggregating counts from each cell and its k nearest neighbors. K was set to 2 for CD26⁺ fibroblast and 1 for FMOD⁺ fibroblast to target ∼10,000 UMI counts per metaCell. No metaCells were generated in myofibroblasts given higher median UMI count. The final count matrix consisted of 250 metaCells from each of the CD26⁺ fibroblast and FMOD⁺ fibroblast populations, and 250 cells randomly sampled from the myofibroblast population. UMI counts were normalized to counts per million (CPM) and transformed by log2(CPM+1). Gene regulatory networks were constructed using ARACNe3, an efficient implementation of the ARACNe-AP algorithm^28^. Two sets of regulators were included, one containing 2,296 mouse transcription factors and cofactors, and one containing 3,197 mouse surface proteins. 50 subnetworks were generated for each set of regulators, consolidated, and pruned at FDR of 0.05. The pruned networks were combined into one network for protein activity inference. Protein activities were inferred from gene expression and gene regulatory network. UMI counts from each fibroblast was CPM-normalized and log2-transformed. Gene expression signatures were obtained by count scaling. The regulatory network was converted to a list of regulons where regulators with <30 targets were removed. The “matrix_narnea” function from PISCES was applied to the gene expression signatures and regulons to infer protein activity measured by proportional enrichment score (PES) implemented in NaRnEA^29^. Master regulators were identified by comparing PES values across fibroblast populations using Kruskal-Walli’s test.

### In silico Perturbation Analysis

Future cell states upon in silico perturbation of a transcription factor were predicted using CellOracle ^30^. The analysis followed the standard CellOracle pipeline with modifications to accommodate batches in the scRNA-seq data. ZsGreen1-negative CD26⁺ and FMOD⁺ fibroblasts were extracted from the integrated analysis and filtered to exclude genes expressed in <10 cells. The top 3,000 highly variable genes were selected using “seurat_v3” method with run ID as the batch key. Highly variable genes were normalized, natural-log transformed, and scaled for downstream analysis. CarDEC embeddings were used to compute neighborhood graph, diffusion map, partition-based graph abstraction (PAGA), and force-directed graph. The root cell for pseudotime calculation was specified as the cell that had ≥95th percentile of *Pi16* expression in CD26⁺ fibroblast and was closest to the middle of the force-directed graph, which corresponded to one end of the CD26⁺ fibroblast trajectory. This choice was used for visualization and does not establish a biological cell of origin. Gene regulatory networks (GRNs) were constructed at each time point and combined to generate consensus GRNs that were robust to batch effect. Specifically, a base GRN was imported at each time point, followed by principal component analysis and k-nearest neighbors (KNN) imputation to construct cluster-specific GRNs. The edges of each GRN were filtered by P-value of 0.001 and the top 10,000 edges were included. A consensus GRN was built for each cluster using GRNs from the 5 time points where an edge was retained if it was present in at least 3 of the 5 time points. The median of the edge coefficients across time points was used as the coefficient for each edge in the consensus GRN. In silico transcription factor (TF) perturbation e.g., for KLF3 and FOXD1, was performed using the preprocessed integrated data and the consensus GRNs. The KNN imputation in the integrated data was modified to use CarDEC embedding instead of principal components to properly identify neighbors in batch corrected latent space. TF expression was set to 0 to simulate knockout and twice the maximum expression to simulate overexpression. Cell transition probability, vector fields, and perturbation scores were calculated and visualized as in the standard pipeline.

### Immunofluorescence

Immunofluorescence was performed on paraffin sections of the aortic root. Sections were deparaffinized in xylene and graded ethanol, followed by antigen retrieval using a Tris-based solution (Vector Laboratories, H3301250). To prevent nonspecific binding, sections were blocked with 10% normal goat serum for 1 hour and incubated overnight at 4C with primary antibodies against PDGFR-α (R&D Systems, AF1062) 1:20, CD68 (Abcam, ab283754), and Cleaved Gasdermin D (Cell Signaling Technology, 10137) at 1:200 concentration, together with the appropriate isotype controls obtained from Thermo-Fisher Scientific. The following day, sections were washed with PBS and incubated with appropriate secondary antibodies for 1 hour at room temperature and mounted with ProLong Glass Antifade Mountant with NucBlue Stain (Thermo-Fisher Scientific, P36983). Imaging was performed on a Nikon Eclipse Ti-S microscope, and images were processed using Image J/Fiji software.

### RNAscope Multiplex Fluorescent Assay v2 (RNA Scope) and RNA-protein Co-detection

RNAscope was performed on formalin-fixed, paraffin-embedded aortic-root sections according to the manufacturer’s instructions (Advanced Cell Diagnostics). Following deparaffinization, target retrieval, hydrogen-peroxide treatment, and protease digestion, sections were hybridized with probes targeting ZsGreen1 (461251-C2), Dpp4 (412751), or Fmod (479412). Signal amplification and fluorophore development were completed using the manufacturer’s amplification reagents and Akoya Biosciences Opal fluorophores. ZsGreen1 was visualized in the Cy5 channel, whereas Dpp4 or Fmod was visualized in the Cy3 channel. To assess the spatial distribution of KLF3 and FOXD1 within aortic-root plaques, ZsGreen1 RNA was detected by RNAscope and KLF3 (Santa Cruz, sc-514500) or FOXD1 (Santa Cruz, sc-293238) protein was detected by immunofluorescence on the same section, following the manufacturer’s combined RNA-protein detection protocol. Slides were mounted with ProLong Glass Antifade Mountant with NucBlue Stain (Thermo Fisher Scientific, P36984). Imaging was performed on a Nikon Eclipse Ti-S microscope, and images were processed using Image J/Fiji software.

### Primary Mouse Aortic Fibroblast Isolation and Culture

Primary aortic fibroblast–enriched populations were isolated from wild-type C57BL/6 mice using an adapted enzymatic dissociation and magnetic cell-selection protocol based on Chen et al^31^. Aortas from two female and two male mice were pooled to generate each biological replicate. Minced aortic tissue was digested for 40 minutes at 37C in DMEM/F12 containing Liberase (1.25mg/mL; Roche, LIBTM-RO), DNase I (0.20 mg/mL; Millipore Sigma, DN25), and hyaluronidase (0.15 mg/mL; Millipore Sigma, H3506) with continuous rotation at 25 rpm using a Fisherbrand tube rotator. The resulting cell suspension was filtered, centrifuged at 4C, and treated with 1× red blood cell lysis buffer (Invitrogen, 00-4333-57). Cells were washed in DPBS containing 0.4% bovine serum albumin and subjected to magnetic depletion of CD45⁺ cells using mouse CD45 MicroBeads (Miltenyi Biotec, 130-052-301). The CD45⁻ fraction was subsequently labeled with a PE-conjugated anti-mouse CD90.2 antibody (BD Biosciences, 553014), followed by Anti-PE MicroBeads (Miltenyi Biotec, 130-048-801) and magnetic positive selection of CD90+ cells. The resulting CD45⁻CD90.2⁺ fraction was used as the fibroblast-enriched population, as we previously described in Bashore et al^12^. Cells were expanded in high-glucose DMEM supplemented with 10% fetal bovine serum, penicillin–streptomycin, 2 mM L-glutamine, and 8 ng/mL fibroblast growth factor 2. Experiments were performed using cells at passages 3–4 unless otherwise indicated.

### Human Aortic Adventitial Fibroblast Culture

Primary human aortic adventitial fibroblasts were obtained from Lonza (CC-7014) and maintained in high-glucose DMEM supplemented with 10% fetal bovine serum, penicillin–streptomycin, 2 mM L-glutamine, and 8 ng/mL fibroblast growth factor 2. Cells were used at passages 4-6 and were cultured at 37°C in a humidified 5% CO₂ incubator.

### siRNA-mediated Gene Silencing

Mouse and human fibroblasts were transfected with TriFECTa Dicer-substrate siRNA duplexes (Integrated DNA Technologies) using Lipofectamine RNAiMAX (Thermo Fisher Scientific, 13778150) in Opti-MEM (Thermo Fisher Scientific, 31985070). A non-targeting DsiRNA was included in every experiment. Lyophilized 2-nmol duplexes were resuspended in 100 μL nuclease-free water to generate 20 μM stocks. Duplexes were combined in equal proportions to a total final siRNA concentration of 40 nM. Mouse *Klf3* knockdown used duplexes Klf3.1 and Klf3.2; human *KLF3* knockdown used KLF3.1, KLF3.2, and KLF3.3. *FOXD1* knockdown used three independent FOXD1 duplexes in mouse and human cells. Cells were plated to reach approximately 60%–70% confluence at transfection. DsiRNA and RNAiMAX were diluted separately in Opti-MEM, combined, incubated for 15 minutes at room temperature, and added to cells in growth medium. Cultures were collected 72 (for Klf3) to 48 (for FoxD1) hours after transfection. Knockdown efficiency was assessed by quantitative PCR and immunoblotting.

### Scratch-wound Migration Assay

Confluent fibroblast monolayers were wounded with a sterile 200-μL pipette tip 48 hours after FOXD1 silencing or 72 hours after KLF3 silencing. Monolayers were washed with PBS to remove detached cells and maintained in medium containing 1% fetal bovine serum. Images of the same wound regions were acquired at 0, 4, and 24 hours. Wound area was measured in ImageJ/Fiji, and percent closure was calculated as [(area at 0 hours − area at the indicated time)/area at 0 hours] × 100.

### RNA Isolation and Reverse Transcription

Cells were rinsed with RNase-free PBS and lysed in TRIzol reagent. Lysates were incubated on ice for 5 minutes. Chloroform was added at one-third the TRIzol volume, samples were mixed by inversion, incubated for 3 minutes at room temperature, and centrifuged at 12,000 × g for 15 minutes at 4C. The aqueous phase was recovered once and applied to Quick-RNA Microprep columns (Zymo Research, R1051). RNA was purified with on-column DNase treatment according to the manufacturer’s instructions and eluted in 15 μL RNase-free water. RNA quantity and purity were assessed by NanoDrop spectrophotometry. Equal RNA input (1–1.5 μg) was reverse transcribed using the High-Capacity cDNA Reverse Transcription Kit (Thermo Fisher Scientific, 4374966).

### Quantitative PCR

Quantitative PCR was performed using TaqMan Fast Advanced Master Mix (Thermo Fisher Scientific, 4444557) on a QuantStudio 7 Flex Real-Time PCR System. Gene-specific TaqMan assays were used to quantify target transcripts. Mouse *Actb, Hprt, and Gapdh* and human ACTB, HPRT, and *GAPDH* were evaluated as endogenous controls. Gapdh and GAPDH, respectively, were selected for normalization based on their stable expression across experimental conditions. Reactions were performed in technical duplicate, and mean Ct values were used to calculate relative gene expression by the 2^−ΔΔ*Ct*^method, normalized to the corresponding non-targeting siRNA or vehicle-control group. Statistical analyses were performed using biological replicates rather than individual technical reactions.

### Immunoblotting

Cells were lysed in 1× RIPA buffer prepared from 10× RIPA Lysis Buffer (Millipore, 20-188) supplemented with PhosSTOP™ phosphatase inhibitor (Roche, 04906837001). Lysates were clarified by centrifugation at 12,000 × *g* for 15 minutes at 4°C, and protein concentrations were determined using the Pierce™ BCA Protein Assay Kit (Thermo Scientific, PI23227). For KLF3 (Santa Cruz, sc-514500) 20 μg of protein per lane was separated by SDS–PAGE using 10% polyacrylamide gels. For FOXD1 (Santa Cruz, sc-293238) analysis, 30 μg of protein per lane was separated using 12% gels. Proteins were transferred to nitrocellulose membranes for 3 minutes using an iBlot™ 3 Western Blot Transfer System. Following blocking and antibody incubation, immunoreactive bands were visualized using Immobilon® ECL Ultra Western HRP Substrate (MilliporeSigma; reagents A and B, CS222619 and CS222618, respectively).

### Statistical Analysis

Statistical analyses were performed using GraphPad Prism, and data are presented as mean ± SEM unless otherwise indicated. For histological analyses, measurements from multiple aortic root sections were averaged for each mouse. Quantification was performed blinded to experimental group. Differences across regression time points were assessed using one-way ANOVA followed by Dunnett’s multiple-comparisons test against baseline, whereas paired two-tailed Student’s t tests were used to compare control and knockdown conditions in the siRNA experiments. Statistical significance was defined as P < 0.05, with sample sizes and statistical tests indicated in the corresponding figure legends.

## Results

### Sustained LDL-C lowering induces progressive plaque remodeling and features of plaque stability

Myh11 CreERT2; Ldlr^−/−^; ROSA26ZsGreen1 mice were used to trace SMC-derived cells and assess changes in plaque cellular composition throughout 20 weeks of sustained LDL-C lowering induced regression (**Figure 1A**). By two weeks, CD68⁺ macrophage content was substantially decreased and continued to decline through later time points (**Figure 1B, C**). This reduction was paralleled by a striking early decrease in cleaved Gasdermin D, a marker of inflammasome-associated pyroptotic cell death, which was nearly absent from lesions by ten weeks **(Figure 1B, D)**. Together, these findings demonstrate early and sustained decreases in inflammatory macrophage accumulation and cell death during regression. Consistent with findings from prior murine models and human regression studies, total lesion area changed minimally during sustained regression (**Figure 1E; Supplemental Fig. 1B**). However, sustained LDL-C lowering was associated with marked compositional remodeling: necrotic core area decreased progressively over time (**Figure 1F; Supplemental Fig. 1C**), while collagen deposition and fibrous cap thickness also increased significantly but more slowly following early inflammatory changes (**Figure 1G, H; Supplemental Fig. 1D, E**). Overall, our prolonged regression protocol revealed major structural remodeling and stabilization, particularly at later time points of LDL-C lowering, rather than significant plaque shrinkage.

**Figure 1.**
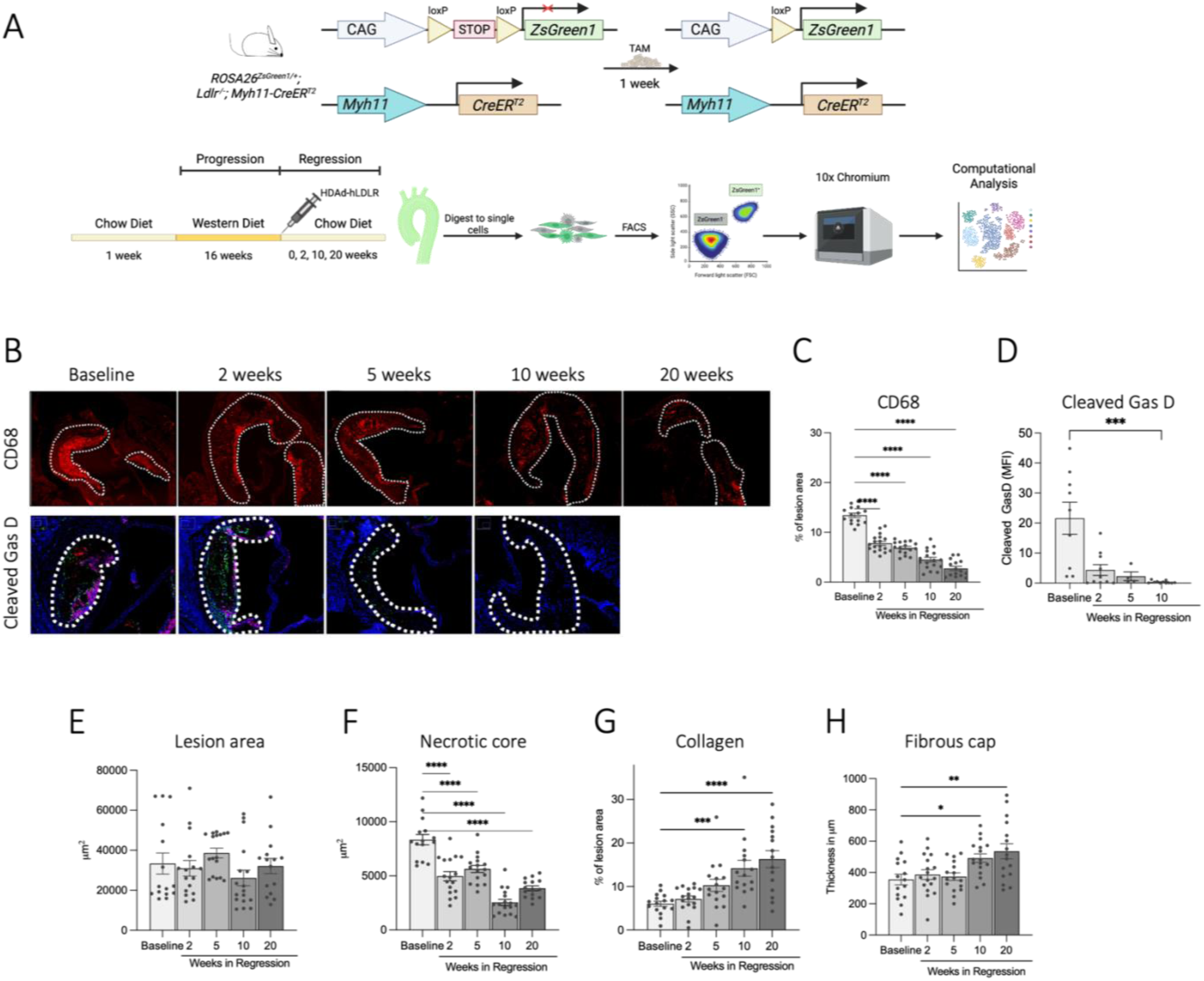
Sustained LDL-C lowering reduces plaque inflammation and promotes features of plaque stability. **A)** Schematic of the *Myh11-CreERT2; Ldlr*−/−; *ROSA26-ZsGreen1* lineage-tracing model and experimental workflow for atherosclerosis progression and regression. Following 16 weeks of Western diet, regression was induced by HDAd-hLDLR administration and transition to chow diet. Aortic-root lesions were analyzed at baseline and after 2, 5, 10, or 20 weeks of regression. **(B)** Representative aortic-root sections stained for CD68 and cleaved gasdermin D during regression. Dashed lines delineate plaque boundaries. **(C)** Quantification of CD68-positive area as a percentage of lesion area. **(D)** Quantification of cleaved gasdermin D mean fluorescence intensity. **(E)** Quantification of total lesion area. **(F)** Quantification of necrotic core area. **(G)** Quantification of collagen-positive area as a percentage of lesion area. **(H)** Quantification of fibrous-cap thickness. Data are presented as mean ± SEM; n=15–17 mice per group. Statistical significance was determined by ordinary one-way ANOVA followed by Dunnett’s multiple-comparisons test, with each regression time point compared with baseline. *P<0.05, **P<0.01, ***P<0.001, and ****P<0.0001.

### Single-cell profiling and SMC lineage tracing reveal early and selective fibroblast subtype expansion during LDL-C lowering induced plaque remodeling

To understand how different cell populations respond to LDL-C lowering, we performed scRNA-seq at baseline and throughout regression (at 0, 2, 5, 10 and 20 weeks of LDL-C lowering), identifying 21 transcriptionally distinct clusters. These included SMC and SMC-derived cells, fibroblasts, macrophages, endothelial cells, and immune populations (**Figure 2A**). Using the Myh11 ZsGreen1 lineage tracing, ZsGreen⁺ cells were classified as SMC and SMC-derived, while ZsGreen⁻ cells represented all non–SMC-derived populations (**Figure 2B**). Although there was a modest increase in the contractile SMC cell population over time, there were almost no changes in the SMC-derived cell subtypes throughout the full course of regression (**Figure 2C–D**). In contrast, specific fibroblast subpopulations showed striking changes: Fmod⁺ and CD26⁺ fibroblast subtypes expanded early by 5 weeks of LDL-C lowering and remained elevated through 20 weeks, whereas the myofibroblast subtype showed no increase and tended to decline (**Figure 2D–E**). These findings suggest that specific fibroblast subtypes, rather than SMC-derived populations, undergo prominent expansion during LDL-C lowering induced remodeling and plaque stabilization.

**Figure 2.**
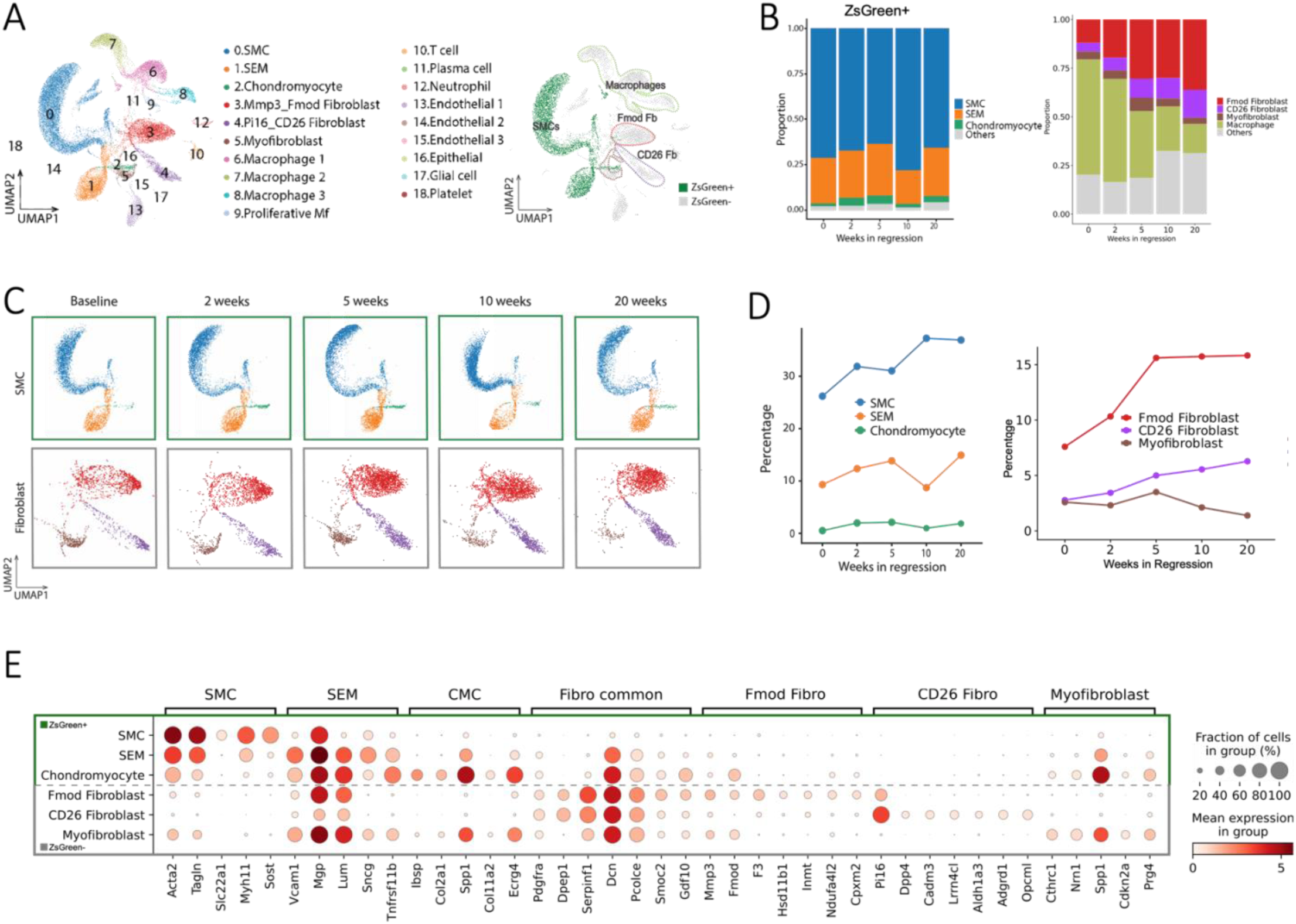
Single-cell profiling and SMC lineage tracing reveal expansion of fibroblast subpopulations during plaque regression. **(A)** UMAP of 21 cell clusters identified by scRNA-seq at baseline and after 2, 5, 10, and 20 weeks of LDL-C lowering, including SMCs, SMC-derived cells, fibroblasts, and immune populations. **(B)** Relative proportions of ZsGreen⁺ SMC-lineage and ZsGreen⁻ non–SMC-lineage cell populations across regression time points. **(C)** UMAP distributions of SMC and fibroblast populations at baseline and throughout regression. **(D)** Percentage of SMC, SEM, and chondromyocyte populations (left) and FMOD⁺, CD26⁺, and myofibroblast populations (right) across regression time points. **(E)** Dot plot showing expression of canonical SMC and fibroblast markers and genes enriched in individual fibroblast populations. Dot size represents the percentage of cells expressing each gene, and color indicates mean gene expression within each population.

To further characterize these fibroblast populations, we examined top differentially expressed genes (DEGs) and canonical fibroblast markers (**Figure 2F; Supplemental Fig. 2A–B**). Several established fibroblast markers, e.g., *Dcn, Pcolce, and Smoc2* were present in all fibroblast populations yet also had high expression in SMC-derived cells. In contrast, Pdgfrα and Serpinf1 were shared across all fibroblast subsets and had minimal expression in SMC-derived cells whereas. *Dpep1* showed robust expression in CD26⁺ and FMOD⁺ fibroblasts but less in myofibroblasts and had no detectable expression in ZsGreen+ SMC-derived populations. This distribution is consistent with recent studies with our collaborators identifying *PDGFRα* and *DPEP1* as robust and complementary markers of adventitial fibroblasts in human and murine arteries^13^. *Mmp3* and *Fmod* were highly expressed in FMOD⁺ fibroblasts but were also expressed in SMC-derived cells, whereas *F3* and at a lower-level *Hsd11b1, Inmt, Ndufa4l2*, and *Cpxm2* were more selectively enriched in Fmod⁺ fibroblasts. *Dpp4* (CD26) was enriched in a subset of *Pi16* expressing fibroblasts, which we labelled CD26⁺ fibroblasts. The cluster annotated as myofibroblasts based on *Acta2, Col1a1*, and *Dcn* expression was enriched for *Spp1* and *Prg4,* yet we did not identify any genes that were selective for myofibroblasts that were not also expressed in SMC-derived populations. These analyses reveal specific fibroblast subpopulations identified in mouse atherosclerotic arteries and their distributional changes in lesions during disease regression. They also highlight the need for more refined molecular signatures to uniquely identify fibroblasts and their subtypes in atherosclerotic lesions

### Spatial validation reveals distinct localization of fibroblast subpopulations in established disease and during plaque regression

To identify the spatial organization of fibroblast subpopulations, we performed RNAscope and immunofluorescence analyses on aortic root sections at 16 weeks of atherosclerosis progression (baseline) and across progressive stages of LDL-C lowering over 10 weeks (**Figure 3**). Fibroblast-associated markers displayed distinct spatial distributions in plaques and during regression. PDGFRα protein was broadly detected within the intima and fibrous cap of the plaque as well as the adventitia, increased in the intima and fibrous cap during regression, and had minimal overlap with ZsGreen⁺ SMC and SMC-derived cells: this supports the expansion of fibroblast populations in lesion intima and fibrous cap in response to sustained lipid lowering (**Figure 3A**). *Dpp4* transcripts, detected by RNAscope had minimal ZsGreen⁺ overlap and became progressively enriched in the intima and within the fibrous cap during prolonged regression (**Figure 3B**). *Fmod* transcripts were detected predominantly in the plaque shoulder and intimal regions, with some overlap with ZsGreen⁺ cells (**Figure 3C**). This overlap is consistent with the scRNA-seq finding that *Fmod* is expressed by both fibroblasts and some SMC-derived cells. Quantification confirmed increased PDGFRα fluorescence intensity and *Dpp4* and *Fmod* RNA signals during regression (**Figure 3A–C**). These findings localize fibroblast-associated gene expression to collagen-rich domains implicated in plaque stabilization during plaque remodeling and support the scRNA-seq finding of increased CD26⁺ and FMOD⁺ fibroblasts in lesions during regression.

**Figure 3.**
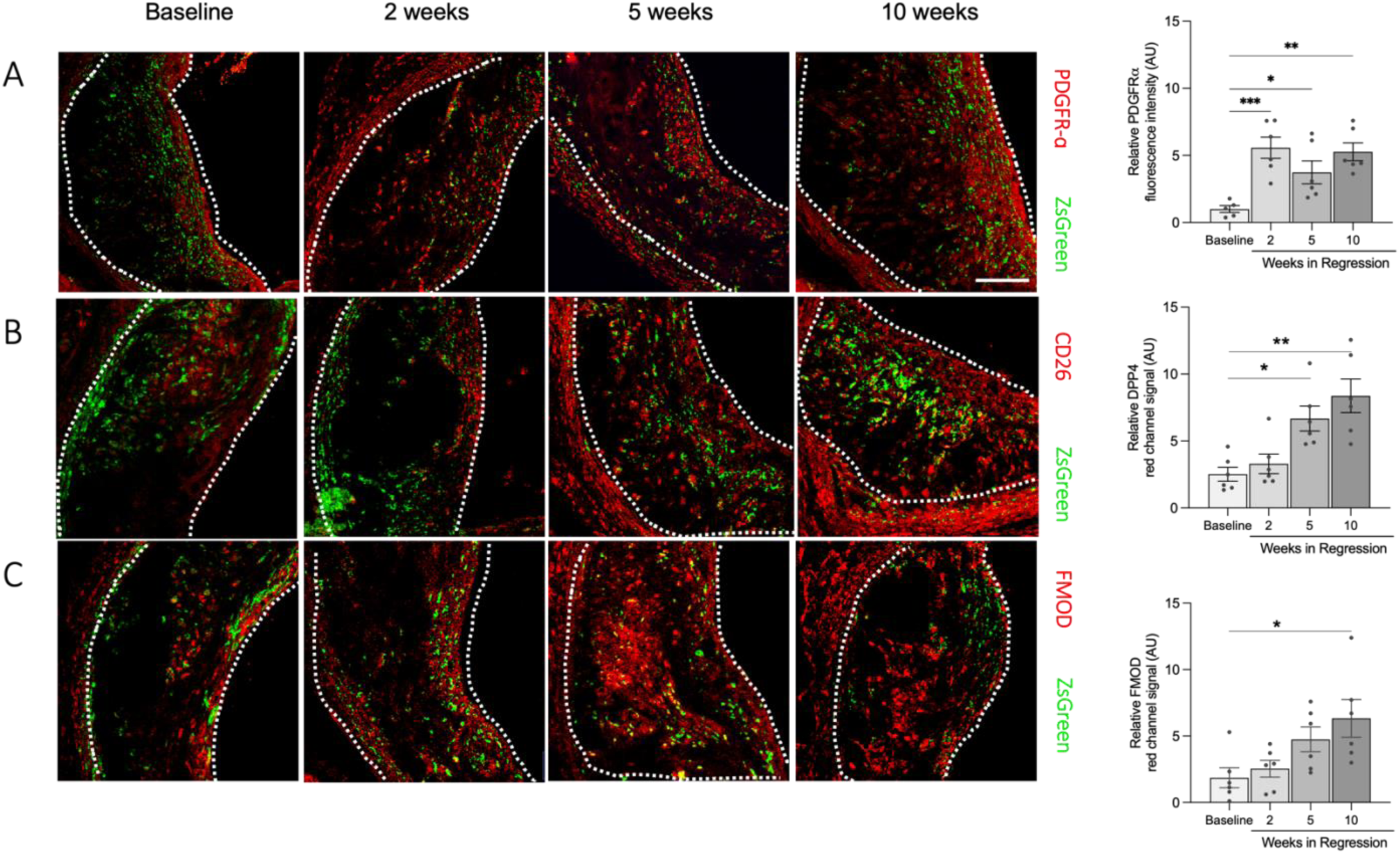
Spatial validation reveals distinct localization of fibroblast-associated markers during plaque regression. **(A)** Representative immunofluorescence images and quantification of PDGFRα in aortic-root lesions at baseline and after 2, 5, or 10 weeks of regression, showing increased signal in the intima and fibrous cap during LDL-C lowering. **(B)** Representative RNAscope images and quantification of Dpp4/CD26 transcripts, showing increased signal in the intima and fibrous cap during regression. **(C)** Representative RNAscope images and quantification of Fmod transcripts, detected predominantly in the plaque shoulder and intimal regions, with partial overlap with ZsGreen⁺ SMC-derived cells. Fibroblast-associated markers are shown in red, ZsGreen⁺ SMC-derived cells in green, and nuclei in blue. ZsGreen was used to assess spatial overlap qualitatively and was not included in the fluorescence quantification. Dashed lines delineate plaque boundaries, and FC indicates the fibrous cap. Right panels show relative fluorescence intensity within the lesion. Scale bars, 50 μm. Data are presented as mean ± SEM; n=6 mice per group. Statistical significance was determined by ordinary one-way ANOVA followed by Dunnett’s multiple-comparisons test, with each regression time point compared with baseline. \**P*<0.05, \*\**P*<0.01, and \*\*\**P*<0.001.

### CD26⁺ and FMOD⁺ fibroblasts engage distinct matrisome programs during plaque regression

The unique temporal pattern of change and spatial distribution of CD26⁺ and FMOD⁺ fibroblasts suggest that they may contribute differently to plaque remodeling. Therefore, we first compared their broader transcriptional programs using GO Biological Process pathway enrichment analysis. This analysis identified enrichment of biological processes related to cell migration, actin filament organization, and cell-substrate adhesion in CD26⁺ , whereas FMOD⁺ fibroblasts were enriched for ECM organization, collagen-related processes, and connective tissue development (**Figure 4B**) We focused next on ECM programs using curated mouse matrisome genes sets developed by Naba et.al., which classify ECM genes into core-matrisome and matrisome-associated categories^23^. CD26⁺ fibroblasts showed higher expression of ECM glycoproteins and matrix regulatory genes, including *Fn1*, *Fbn1*, *Lama4*, *Vcan*, and *Dpt*. In contrast, FMOD⁺ fibroblasts preferentially expressed genes involved in collagen, proteoglycan, and elastic-matrix organization, including *Col1a1*, *Col4a1*, *Col6a3*, *Bgn*, *Lum*, *Eln*, and *Fmod* (**Figure 4A**). Consistent with these gene-level differences, gene-set enrichment analysis showed that CD26⁺ fibroblasts were enriched for ECM glycoprotein, ECM-affiliated, and matrisome-regulatory programs, whereas FMOD⁺ fibroblasts were enriched for collagen and proteoglycan programs (**Figure 4C**). Together, these findings support distinct but complementary roles for CD26⁺ and FMOD⁺ fibroblasts in ECM reorganization during LDL-C lowering induced plaque remodeling.

**Figure 4.**
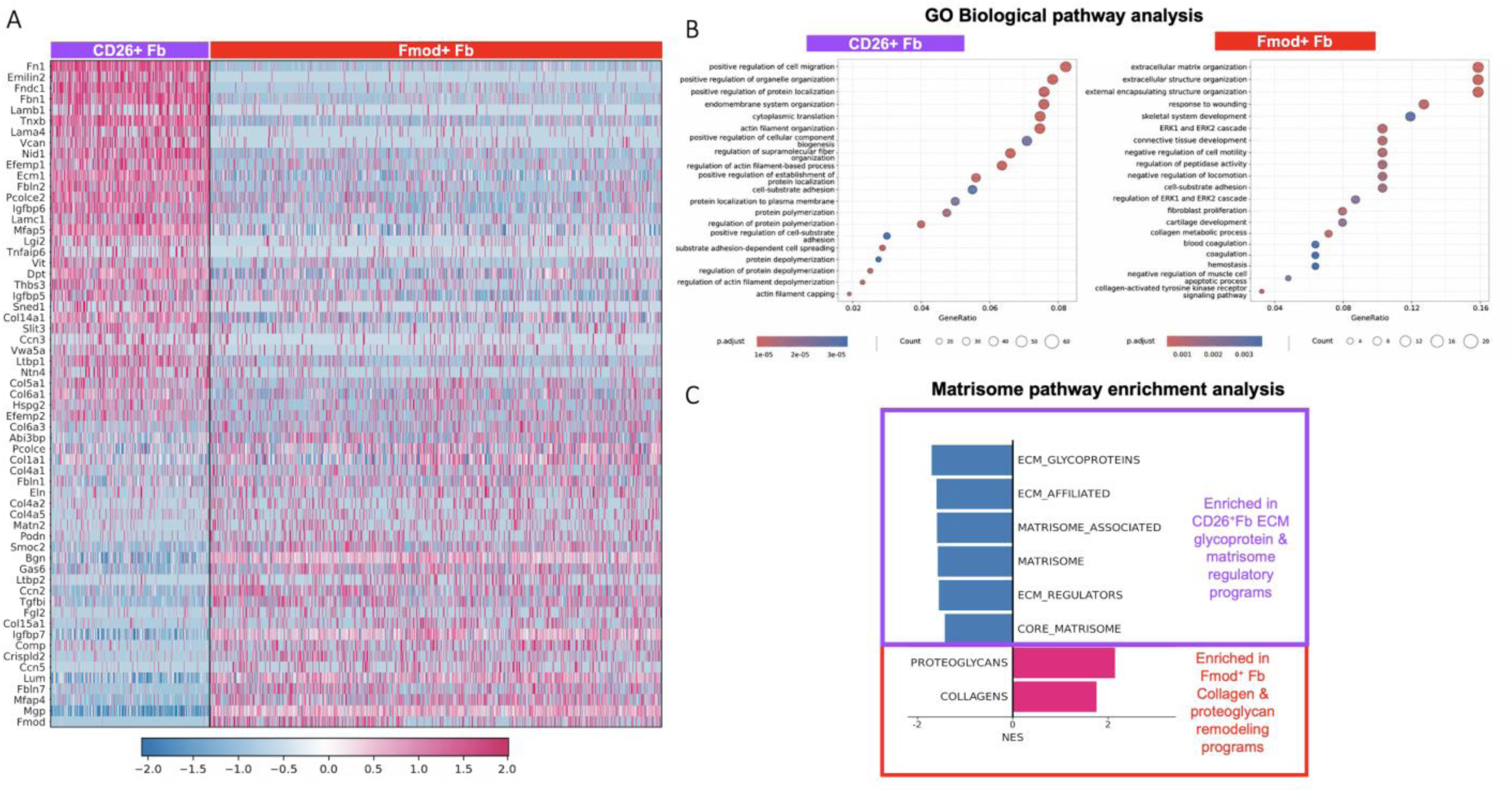
CD26⁺ and FMOD⁺ fibroblasts exhibit distinct cellular and matrisome programs during plaque regression. **(A)** Heatmap showing scaled expression of representative matrisome genes across individual CD26⁺ and FMOD⁺ fibroblasts identified by scRNA-seq. Columns represent individual cells grouped by fibroblast population, and rows represent selected matrisome genes. The upper gene block is enriched in CD26⁺ fibroblasts and includes ECM glycoproteins and matrisome regulators, whereas the lower block is enriched in FMOD⁺ fibroblasts and includes genes associated with collagen, proteoglycan, and elastic-matrix organization. **(B)** GO Biological Process enrichment analysis of differentially expressed genes between CD26⁺ and FMOD⁺ fibroblasts. CD26⁺ fibroblasts showed enrichment of pathways associated with positive regulation of cell migration, actin filament organization, and cell–substrate adhesion, whereas FMOD⁺ fibroblasts were enriched for ECM organization, collagen metabolism, fibroblast proliferation, and negative regulation of cell motility. Dot size represents the number of genes associated with each pathway, dot color indicates the adjusted P-value, and the x-axis shows the gene ratio. **(C)** Complementing the matrisome gene-expression patterns shown in panel A, gene-set enrichment analysis of curated mouse matrisome gene sets from MSigDB and the Matrisome Project further characterized the distinct ECM programs of CD26⁺ and FMOD⁺ fibroblasts. A total of 5,094 genes expressed in at least 10% of CD26⁺ or FMOD⁺ fibroblasts were ranked by log₂ fold change between the two populations. Negative normalized enrichment scores indicate enrichment in CD26⁺ fibroblasts, whereas positive scores indicate enrichment in FMOD⁺ fibroblasts. CD26⁺ fibroblasts were enriched for ECM glycoprotein, ECM-affiliated, and matrisome-regulatory programs, whereas FMOD⁺ fibroblasts were enriched for collagen and proteoglycan programs

### Regulatory network inference identifies KLF3 and FOXD1 as candidate regulators of fibroblast state specialization during LDL-C lowering

To identify transcriptional regulators associated with fibroblast subtype identity during sustained LDL-C lowering, we applied ARACNe network reconstruction followed by NaRnEA protein activity inference to fibroblast subpopulations identified by scRNA-seq (**Figure 5**). This analysis revealed distinct regulatory signatures across CD26⁺ fibroblasts, FMOD⁺ fibroblasts, and myofibroblasts during plaque regression (**Figure 5A**). The predicted activity of KLF3, a regulator of adipose stromal cell differentiation and suppressor of innate immunity^32, 33^, was selectively elevated in CD26⁺ fibroblasts and reduced in FMOD⁺ fibroblasts and myofibroblasts (**Figure 5A, B**). In contrast, the predicted activity of FOXD1, a known regulator of kidney stromal cells and kidney development as well as kidney and lung fibrosis^34–36^, was prominently and selectively activated in FMOD⁺ fibroblasts (**Figure 5A, C**). Notably, the inferred protein activity patterns were much more clearly segregated by fibroblast subtypes than their transcript expression alone, suggesting CD26⁺ and FMOD⁺ fibroblast subtype-specific activation of KLF3- and FOXD1-regulated networks, respectively. Immunofluorescence staining further detected KLF3 and FOXD1 in aortic root lesions at baseline and after 20 weeks of regression (**Figure 5D, E**). Together, these findings identified KLF3 and FOXD1 as candidate regulators associated with CD26⁺ and FMOD⁺ fibroblast transcriptional programs during plaque remodeling after LDL-C lowering.

**Figure 5.**
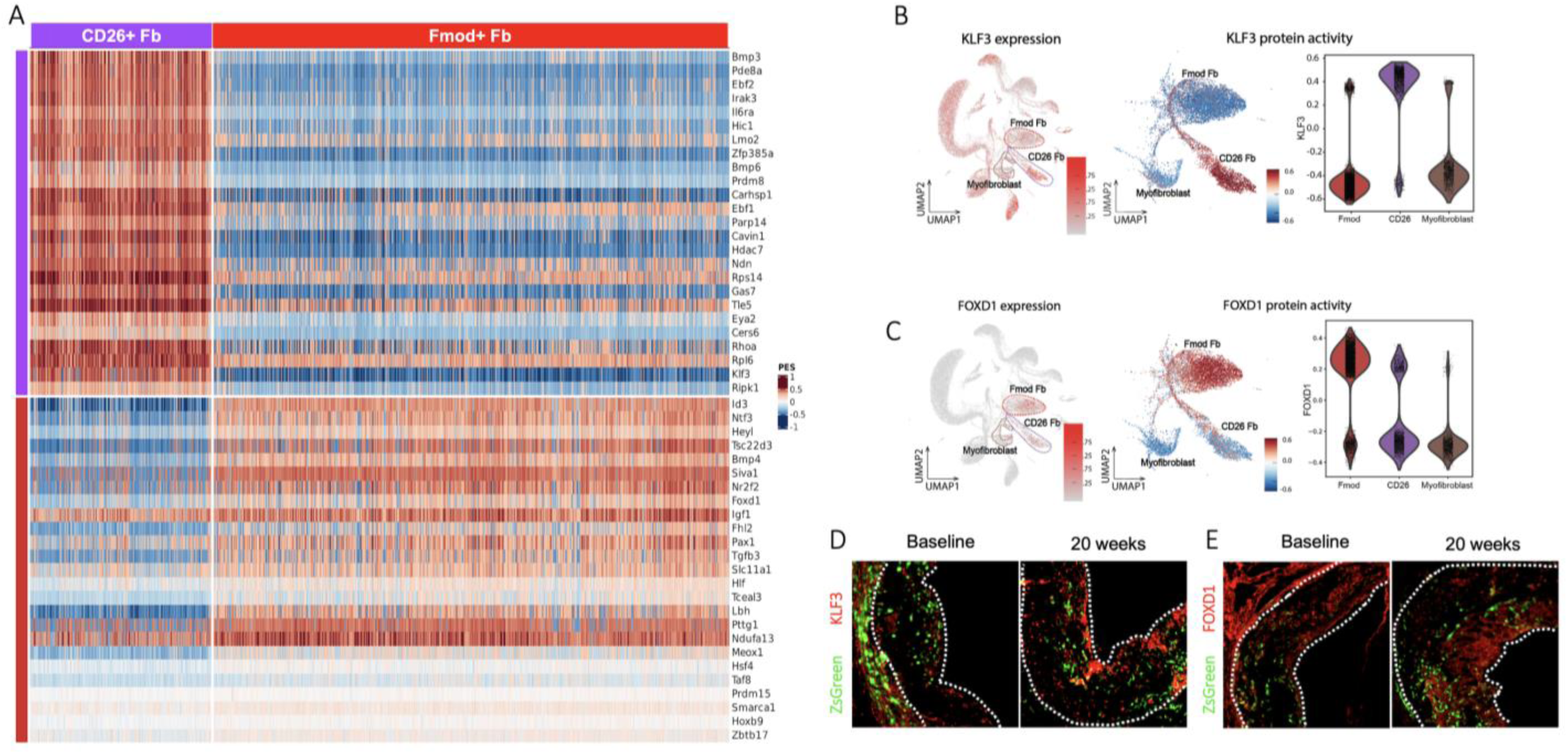
Regulatory network inference identifies distinct KLF3- and FOXD1-associated programs in fibroblast populations during plaque regression. **(A)** Heatmap showing inferred transcriptional regulator activity across individual CD26⁺ and FMOD⁺ fibroblasts identified by scRNA-seq. A fibroblast-specific regulatory network was reconstructed using ARACNe, followed by NaRnEA regulon enrichment analysis to infer regulator protein activity. Columns represent individual cells grouped by fibroblast population, and rows represent candidate transcriptional regulators. The upper regulator block is enriched in CD26⁺ fibroblasts and includes KLF3, whereas the lower block is enriched in FMOD⁺ fibroblasts and includes FOXD1. **(B)** UMAP feature plots showing Klf3 transcript expression and inferred KLF3 protein activity, with violin plots comparing inferred activity across CD26⁺, FMOD⁺, and myofibroblast populations. **(C)** Corresponding UMAP feature plots and violin plots showing Foxd1 transcript expression and inferred FOXD1 protein activity across fibroblast populations. Color scales indicate relative transcript expression or NaRnEA protein enrichment score. **(D)** Representative RNAscope–immunofluorescence images showing KLF3 protein (red) and ZsGreen1 RNA (green) in aortic-root lesions at baseline and after 20 weeks of regression, with KLF3 signal detected within the plaque intima, particularly near the fibrous cap**. (E)** Corresponding images showing FOXD1 protein (red) and ZsGreen1 RNA (green), with FOXD1 signal detected in the vessel wall and plaque intima. Dashed lines delineate plaque boundaries.

### KLF3 supports the CD26⁺ fibroblast matrix-modulating functions and restrains inflammatory activation

To probe the functional role of KLF3 in fibroblast specialization, we used pseudotime analysis and CellOracle perturbation modeling of our *ex vivo* scRNA-seq regression data as well as independent experimental KLF3 silencing in primary mouse and human aortic fibroblasts (**Figure 6**). Pseudotime analysis identified a transcriptional continuum between CD26⁺ and FMOD⁺ fibroblasts, consistent with dynamic remodeling of these sub-populations during regression (**Figure 6A**). CellOracle predicted that *Klf3* loss shifts fibroblast-state dynamics away from the CD26⁺ region towards that associated with the FMOD⁺ region, whereas simulated *Klf3* overexpression favors the CD26⁺ associated program. We next tested whether experimental KLF3 loss alters the molecular and functional properties of cultured fibroblasts. In primary mouse aortic fibroblasts, siRNA-mediated silencing reduced *Klf3* mRNA and KLF3 protein abundance (**Figure 6B**). *Klf3* silencing decreased expression of the CD26⁺ fibroblast marker *Dpp4* without significantly altering *Pdgfra*. Expression of contractile genes *Acta2* and *Myh11,* proliferation genes *Ki67* and *Ccnd1*, and the collagens *Col1a1*, *Col1a2*, and *Col3a1* were also modestly reduced. In parallel, *Klf3* silencing increased inflammatory chemokines and mediators including *Cxcl16, Il6, Ccl2*, and *Lgals3* (**Figure 6B**), suggesting that suppression of KLF3 shifts fibroblasts away from matrix-producing functions while promoting inflammatory gene expression. Consistent with these molecular effects, *Klf3*-silenced mouse fibroblasts showed impaired wound closure in scratch assays (**Figure 6C**). In human fibroblasts (**Figure 6D**), KLF3 silencing similarly reduced *DPP4*, *ACTA2*, *MYH11*, *MKI67*, *CCND1*, *COL1A1*, *COL1A2*, and *COL3A1* expression, whereas *PDGFRA* remained unchanged, and showed impaired wound closure (**Figure 6E**). Together, these findings support a conserved role for KLF3 in regulating matrix-associated gene expression and scratch closure in cultured mouse and human fibroblasts, with additional effects on inflammatory and chemotactic genes in mouse fibroblasts.

**Figure 6.**
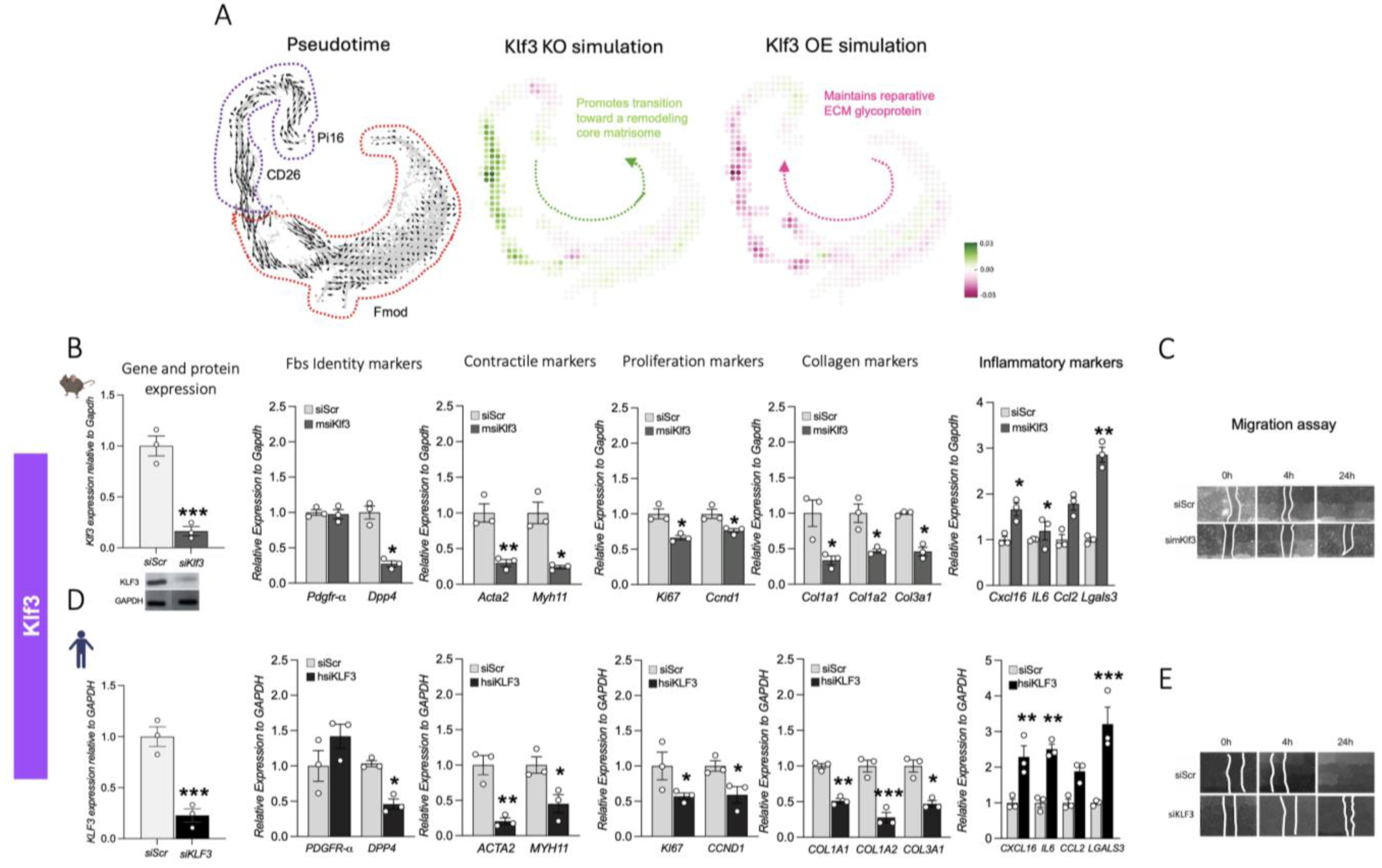
KLF3 supports the CD26⁺ fibroblast program and regulates matrix-associated and inflammatory gene expression. **(A)** Pseudotime trajectory and CellOracle perturbation analysis of CD26⁺ and FMOD⁺ fibroblasts. The pseudotime plot shows the inferred transcriptional continuum between these populations. CellOracle simulations show predicted changes in fibroblast-state dynamics following *Klf3* knockout or overexpression. Simulated *Klf3* loss shifts predicted dynamics toward the FMOD⁺ fibroblast-associated program, whereas *Klf3* overexpression favors the CD26⁺ fibroblast-associated program. Colors indicate predicted perturbation scores, and arrows indicate the direction of inferred cell-state transitions. **(B)** Validation of *Klf3* silencing in primary mouse aortic fibroblasts by quantitative PCR and Western blotting, followed by expression analysis of fibroblast markers (*Pdgfra* and *Dpp4*), contractile genes (*Acta2* and *Myh11*), proliferation-associated genes (*Mki67* and *Ccnd1*), collagen genes (*Col1a1, Col1a2,* and *Col3a1*), and inflammatory mediators (*Cxcl16, Il6, Ccl2,* and *Lgals3*). **(C)** Representative scratch-wound images of control and *Klf3*-silenced mouse fibroblasts at 0, 4, and 24 hours after injury. **(D)** Validation of *KLF3* silencing and expression analysis of *PDGFRA, DPP4*, *ACTA2, MYH11, MKI67, CCND1, COL1A1, COL1A2*, and *COL3A1* in primary human aortic fibroblasts. **(E)** Representative scratch-wound images of control and *KLF3*-silenced human fibroblasts at 0, 4, and 24 hours after injury. Gene expression was normalized to *GAPDH* and expressed relative to the corresponding control. *GAPDH* was also used as the protein-loading control. Data are presented as mean ± SEM; n=3 independent experiments. For each gene, control and *KLF3*-silenced conditions were compared using a paired two-tailed Student’s t test. \**P*<0.05, \*\**P*<0.01, and \*\*\**P*<0.001.

### FOXD1 supports the FMOD⁺ fibroblast program and matrix-producing functions

*In silico* CellOracle perturbation (**Figure 7A**) predicted that *Foxd1* loss favors the CD26⁺ fibroblast-associated ECM-glycoprotein program, whereas simulated *Foxd1* overexpression shifts fibroblast-state dynamics toward the FMOD⁺ fibroblast core-matrisome program. In primary mouse fibroblasts, siRNA-mediated silencing reduced FOXD1 expression (**Figure 7B**) and markedly suppressed both *Fmod* and the canonical fibroblast marker *Pdgfra*. This was accompanied by substantial reduction of *Acta2* and *Myh11* expression as well as *Mki67* and *Ccnd1*, and marked reductions in *Col1a1*, *Col1a2*, and *Col3a1*(**Figure 7B**), as well as impaired wound closure (**Figure 7C**). In primary human fibroblasts, FOXD1 silencing similarly and markedly reduced *PDGFRA*, *FMOD*, *ACTA2*, *MYH11*, *MKI67*, *CCND1*, *COL1A1*, *COL1A2*, and *COL3A1* expression **(Figure 7D)** and impaired wound closure (**Figure 7E**). Together, these findings support a conserved role for FOXD1 in maintaining fundamental fibroblast identity as well as in collagen and matrix-producing, proliferative, and migratory functions.

**Figure 7.**
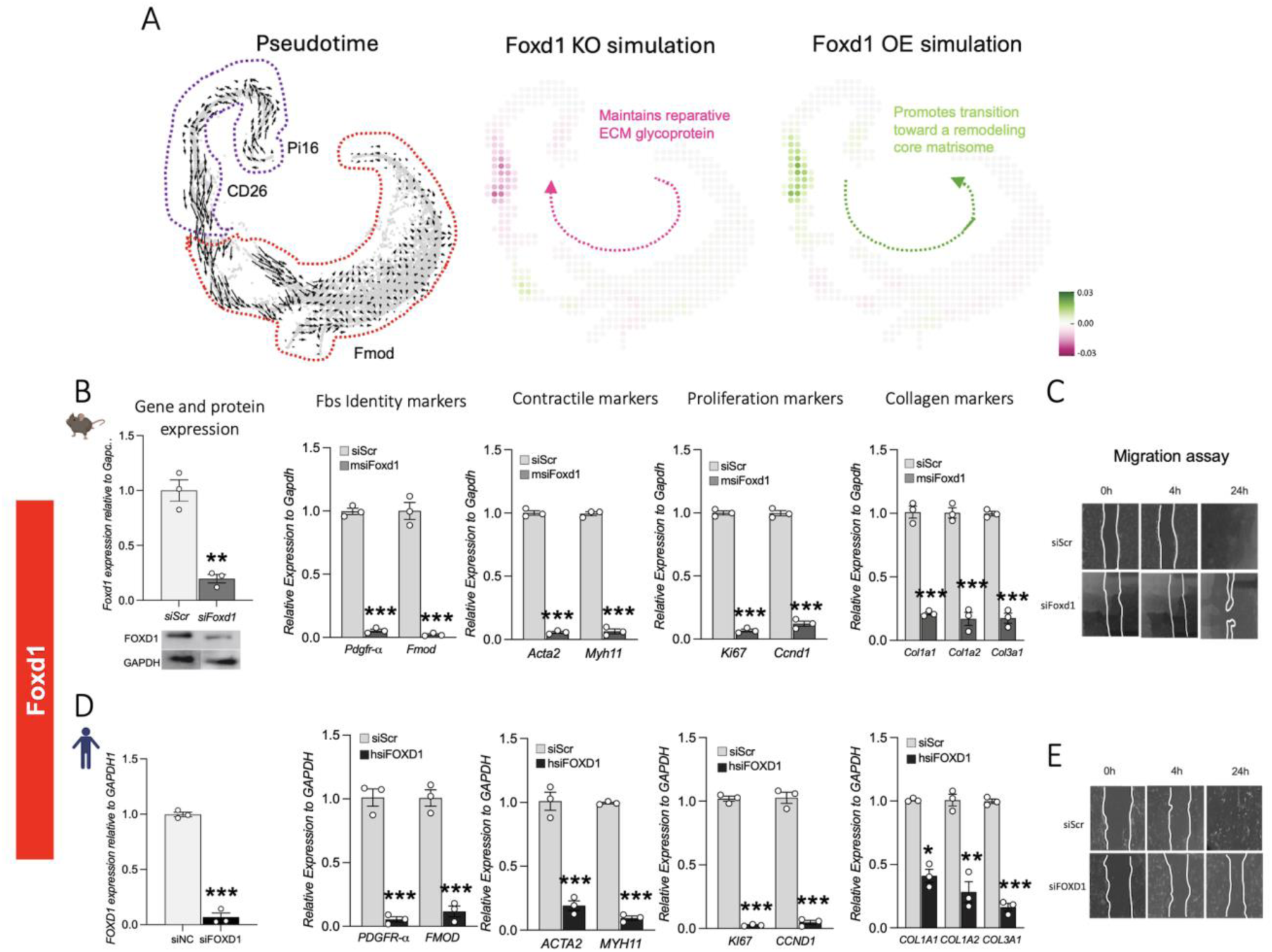
FOXD1 supports the FMOD⁺ fibroblast program and regulates matrix-associated gene expression. **(A)** Pseudotime trajectory and CellOracle perturbation analysis of CD26⁺ and FMOD⁺ fibroblasts. The pseudotime plot shows the inferred transcriptional continuum between these populations. CellOracle simulations show predicted changes in fibroblast-state dynamics following *Foxd1* knockout or overexpression. Simulated *Foxd1* loss favors the CD26⁺ fibroblast-associated program, whereas *Foxd1* overexpression shifts predicted dynamics toward the FMOD⁺ fibroblast-associated program. Colors indicate predicted perturbation scores, and arrows indicate the direction of inferred cell-state transitions. **(B)** Validation of *Foxd1* silencing in primary mouse aortic fibroblasts by quantitative PCR and Western blotting, followed by expression analysis of fibroblast markers (*Pdgfra* and *Fmod*), contractile genes (*Acta2* and *Myh11*), proliferation-associated genes (*Mki67* and *Ccnd1*), and collagen genes (*Col1a1, Col1a2,* and *Col3a1*). **(C)** Representative scratch-wound images of control and *Foxd1*-silenced mouse fibroblasts at 0, 4, and 24 hours after injury. **(D)** Validation of FOXD1 silencing and expression analysis of *PDGFRA, FMOD, ACTA2, MYH11, MKI67, CCND1, COL1A1, COL1A2,* and *COL3A1* in primary human aortic fibroblasts. **(E)** Representative scratch-wound images of control and *FOXD1*-silenced human fibroblasts at 0, 4, and 24 hours after injury. Gene expression was normalized to *GAPDH* and expressed relative to the corresponding control. *GAPDH* was also used as the protein-loading control. Data are presented as mean ± SEM; n=3 independent experiments. For each gene, control and *FOXD1*-silenced conditions were compared using a paired two-tailed Student’s t test. \**P*<0.05, \*\**P*<0.01, and \*\*\**P*<0.001.

### Human genetic analyses identify associations of KLF3 and FOXD1 to coronary artery disease

To examine the human relevance of our findings, we assessed associations between genetic variation in KLF3 and FOXD1 and coronary artery disease (CAD). In the Million Veteran Program, the KLF3-associated variant rs16994723 showed a suggestive association with CAD (P=5.61×10⁻⁴). Gene-based burden testing identified a nominal association between FOXD1 variation and CAD (P=0.018).

## Discussion

We combined single-cell profiling and lineage tracing to provide the first longitudinal analysis of vascular stromal cell and lesion structural responses during sustained lipid lowering in mice with established atherosclerosis. To our surprise, there were minimal change in SMC-derived cells, whereas CD26⁺ and FMOD⁺ fibroblast subtypes increased markedly as plaques developed thicker fibrous caps, increased collagen content, and smaller necrotic cores. Fb subtype expression profiles distinguished ECM glycoprotein and regulatory programs in CD26⁺ fibroblasts from collagen and proteoglycan programs in FMOD⁺ fibroblasts, suggesting distinct and potentially complementary functions during lesion remodeling. Regulatory network analysis and CellOracle predictions nominated KLF3 and FOXD1 as master regulators of CD26⁺ and FMOD⁺ fibroblast subtypes, respectively. Silencing of either factor had distinct and overlapping effects on fibroblast identity, reductions in matrix-associated gene expression and impaired scratch closure in mouse and human aortic fibroblasts. KLF3 silencing also selectively increased inflammatory and chemokine genes in fibroblasts of both species. These findings reveal unique fibroblast subtype-specific responses in lesions during LDL-C lowering and identify KLF3 and FOXD1 as candidate transcriptional regulators of fibroblast matrix-associated and inflammatory functions directly relevant to plaque remodeling and stabilization after LDL-C lowering.

Most work to date has focused on SMC-derived cells as the stromal cell type of relevance in atherosclerosis progression and lesion stability^8–10, 37^, although recent work suggests the presence and potential role of fibroblasts in mouse and human atherosclerotic lesions^11–14^. In mouse atherosclerosis, adventitial fibroblasts undergo changes in their transcriptional programs during disease progression, with SERPINH1 contributing to fibroblast migration^14^, suggesting functions beyond structural support within the adventitia. Our previous work identified a distinct CD26⁺ fibroblast population that expands within atherosclerotic lesions and localizes to the intima and fibrous cap with minimal overlap with SMC-derived cells^12^. Consistent with fibroblasts representing an additional stromal population within plaques, lineage-tracing studies demonstrated that most plaque fibroblasts in intima and fibrous cap originate from adventitial *Pdgfra*-lineage cells and largely maintain fibroblast identity^11^. Liu et al. linked macrophage TREM2–PDGFB signaling to fibroblast accumulation and fibrous-cap remodeling, providing evidence that fibroblasts respond to signals within the plaque microenvironment and may contribute to plaque stability^38^. The relevance of these findings extends to human arteries, where distinct fibroblast populations have been linked to atherosclerotic plaque characteristics^13^, and their transcriptional programs show enrichment for vascular disease-associated genetic signals^39^. These studies support a role for fibroblasts in plaque composition and stability, but how specific fibroblast populations respond to sustained LDL-C lowering and contribute to plaque stabilization and reducing residual CVD risk remains poorly understood.

Here, the increase in fibroblast subtypes, but not SMC-derived cell types, during sustained lipid lowering is a striking contrast to findings during atherosclerosis progression^11–14^. Our previous work identified several fibroblast populations, including a CD26⁺ fibroblast subset, in the adventitia and lesion intima during disease progression; however, SMC-derived cells but not fibroblasts are the cell types that expand most markedly in lesions during hyperlipidemia-induced disease development^12^. During regression, increased PDGFRα and CD26 signals in the intima and fibrous cap localize fibroblast-associated changes to regions undergoing plaque remodeling – not simply to the adventitia, their residence in health vascular homeostasis. Within the fibroblast subtypes, CD26⁺ cells showed higher expression of genes encoding ECM glycoproteins and matrix regulators, whereas FMOD ⁺ cells showed higher expression of collagen and proteoglycan genes. These differences suggest that fibroblast involvement in plaque regression encompasses regulation of the extracellular environment as well as structural matrix production. Thus, alongside established contributions from SMC-derived cells, fibroblasts – through their coordinated subtype-specific functions-may play especially important roles in plaque remodeling and stabilization following LDL-C lowering^8, 9^.

Given population expansion and differences in molecular and predicted functions, we probed how CD26⁺ and FMOD⁺ arterial fibroblast subtypes and programs are regulated. We used ARACNe to reconstruct regulatory networks and NaRnEA to estimate transcription factor activity from predicted target-gene expression^28, 29^, identifying enriched KLF3 activity in CD26⁺ fibroblasts and FOXD1 activity in FMOD⁺ fibroblasts. CellOracle loss-of-function simulations for Klf3 and Foxd1 predicted shifts respectively in CD26⁺ and FMOD⁺ fibroblast transcriptional and cell state phenotypes^30^, suggesting that these factors help maintain specific fibroblast subtype gene programs. Furthermore, published work provides distinct biological rationales to prioritize each candidate. KLF3 regulation of stromal cell-to-adipocyte and adipose maturation as well as its repression of inflammatory genes raised the possibility that its activity influences both inflammatory as well as stromal matrix-associated programs^32, 33, 40, 41^. Studies linking *Foxd1*-lineage stromal cells to matrix-producing populations in kidney and lung development, injury, and fibrosis provide a context for examining FOXD1 regulatory function in vascular fibroblasts^34, 35^. Together, the activity patterns, perturbation predictions, and published biology suggest that both KLF3 and FOXD1 may influence specific arterial fibroblast functions rather than simply be transcriptional markers of fibroblast subtype identities.

KLF3 is a zinc-finger transcription factor that binds CACCC and GC-rich DNA sequences and represses target genes by recruiting C-terminal binding proteins (CtBP1/2) and associated chromatin-modifying corepressor complexes^42–46^. Its repressive activity is enhanced by HIPK2-mediated phosphorylation, which promotes DNA binding and transcriptional repression^45^, and by SUMOylation, which further strengthens transcriptional repression46. It’s established repressor function provides a mechanistic context for its effects in arterial fibroblasts. Previous studies showed that KLF3 suppresses LPS-induced, NF-κB–driven inflammation through repression of *Rela*, which encodes p65, as well as directly repressing *Lgals3*^33, 40^. We found increased *Lgals3* and *Cxcl16* expression, with smaller increases in *Ccl2* and *Il6*, after knockdown of *Klf3* in cultured aortic fibroblasts. This is consistent with a novel function for KLF3 to suppress fibroblast and lesion inflammatory programs during LDL-C lowering mediated regression. KLF3 depletion in cultured fibroblasts also reduced proliferation- and collagen associated transcripts and wound scratch closure, suggesting that KLF3 supports fibroblast matrix remolding in lesions while suppressing inflammatory gene expression. Wound-healing studies show that inflammatory signals influence both repair and fibrosis, providing a remodeling context for integrated Klf3 functions in atherosclerosis regression^15^. The mixed changes in ECM-associated transcripts similarly argue against a uniformly profibrotic or antifibrotic function. KLF3 may instead help establish conditions favorable for remodeling through selective effects on, matrix-associated transcription, inflammatory and chemokine gene expression and potentially fibroblast suppression of macrophage recruitment.

FOXD1 is a forkhead box transcription factor involved in stromal development and extracellular matrix regulation^36, 47^. In the embryonic kidney, FOXD1 represses *Dcn*, which encodes the matrix proteoglycan decorin^47^. Previous studies showed that cells derived from *Foxd1*-expressing progenitors contribute to matrix-producing populations in kidney and lung fibrosis^34, 35^. Whereas those studies established cellular origin, our siRNA-mediated knockdown experiments showed that reducing FOXD1 expression in cultured aortic fibroblasts markedly decreased fibroblast identity and proliferation markers as well as structural collagen transcripts. Consistent with enriched FOXD1 activity in FMOD⁺ fibroblasts and their collagen-and proteoglycan-associated expression profile, these data support a potential role for FOXD1 in maintaining arterial fibroblast identity and expansion as well as collagen production and matrix integrity during arterial lesion remodeling. Fibromodulin’s established role in collagen fibril assembly and organization^48, 49^ as well as in arterial remodeling and experimental atherosclerosis^50–54^ reinforces the potential role of FMOD⁺ fibroblasts and FoxD1 in arterial matrix remodeling during regression. Although matrix production is typically increased in pathological fibrosis, it is also critical for structural support within the atherosclerotic fibrous cap^4, 9^. FOXD1 activity may thus support fibroblast functions directly relevant to fibrous cap remodeling.

Findings in human aortic fibroblasts support conservation of specific regulatory functions in humans as well as mice. KLF3 knockdown reduced *DPP4* and selected collagen transcripts while leaving *PDGFRA* expression unchanged, whereas FOXD1 knockdown reduced *PDGFRA*, *DPP4* and *FMOD*, as well as selected collagen transcripts. Both interventions also impaired scratch closure, and KLF3 knockdown increased selected inflammatory genes in human fibroblasts. Our common variant interrogation analyses in the Million Veteran Program GWAS^55^ and rare variant gene burden testing in our whole exome/whole genome sequencing coronary artery disease datasets provides suggestive evidence for their association with CAD and MI. The common-variant and gene burden analyses address different genetic hypotheses, and neither proves an effect through fibroblasts on plaque stabilization. However, these human fibroblast studies and genetic data provide strong premise for additional human translational and experimental studies to evaluate fibroblast regulation in stabilizing human atherosclerotic plaques.

Our work raises several questions. Are fibroblast KLF3 or FOXD1 activity, and CD26⁺ or FMOD⁺ fibroblasts, required for fibroblast subtype specific functions and plaque remodeling *in vivo* during LDL-C lowering, or in atherosclerosis progression – these questions are the focus of ongoing studies in complex mouse models. FMOD is also expressed in some phenotypically modulated SMC-derived cell that acquire fibroblast-like, matrix-producing features^8^ emphasizing the need for more specific fibroblast subtype lineage markers^11^ for *in vivo* and translational studies. For KLF3, *in vitro* and *in vivo* studies are required to test whether fibroblast secreted CCL2, CXCL16, or LGALS3 contributes to altered macrophage recruitment and behavior in atherosclerosis progression and remodeling. Defining the direct transcriptional targets of KLF3 and FOXD1 and testing whether regulation by TGFβ or PDGF differentially regulate these pathways would further distinguish primary regulatory effects from secondary changes in cell state in response to environmental and niche queues. KLF4, an established regulator of SMC phenotypic modulation^37^, is a candidate that may drive interactions among KLF-family factors in vascular fibroblasts; cooperation or compensation with KLF3 in stromal cells remains untested. Overall, such studies would establish how CD26⁺ and FMOD⁺ fibroblast subtype transcriptional regulation and communication with neighboring cells contribute to plaque remodeling and stabilization.

This work identifies fibroblast subtype responses as a significant component of the cellular and structural remodeling that stabilizes atherosclerotic plaques following sustained lipid lowering. The regulatory and matrix differences between CD26⁺ and FMOD⁺ fibroblasts, together with the effects of KLF3 and FOXD1 knockdown, suggest that fibroblast matrix-associated and inflammatory functions may help determine the course of plaque remodeling after lipid lowering.

## Acknowledgements

We thank the JP Sulzberger Columbia Genome Center for single-cell sequencing support, the Columbia Stem Cell Initiative Flow Cytometry Core Facility for cell sorting, and the Columbia University Molecular Pathology Shared Resource for histological staining. We also thank the Confocal and Specialized Microscopy Shared Resource at the Herbert Irving Comprehensive Cancer Center, Columbia University, for microscope training and access to imaging equipment. This shared resource is supported in part by NIH/NCI Cancer Center Support Grant P30CA013696.

