## Supplemental material for "Sustained LDL-C Lowering Reveals Selective Expansion of Fibroblast Subpopulations with Distinct Regulatory and Matrix Programs During Atherosclerosis Remodeling"

**Supplemental Data**

**Supplemental Table A1:** Filtering parameters applied to mouse scRNA-seq data. The minimum and the maximum number of genes, and the maximum of UMIs applied are shown for each sequencing run. A maximum of 10% reads mapped to mitochondrial genes are allowed in each cell.

| Regression (weeks) | SMC lineage | Min. #genes | Max. #genes | Max. #UMIs | #cells | #cells after filtering (%) |
|---|---|---|---|---|---|---|
| 0 | ZsGreen+ | 200 | 4,000 | 20,000 | 6,607 | 4,711 (71.3) |
| | ZsGreen- | | 3,500 | 15,000 | 8,375 | 8,077 (96.4) |
| 2 | ZsGreen+ | | 4,500 | 20,000 | 7,033 | 5,579 (79.3) |
| | ZsGreen- | | 4,000 | 20,000 | 6,500 | 6,196 (95.3) |
| 5 | ZsGreen+ | | 4,000 | 20,000 | 8,394 | 5,570 (66.4) |
| | ZsGreen- | | 3,500 | 15,000 | 6,309 | 5,853 (92.8) |
| 10 | ZsGreen+ | | 4,000 | 20,000 | 6,674 | 5,099 (76.4) |
| | ZsGreen- | | 3,000 | 10,000 | 5,881 | 5,603 (95.3) |
| 20 | ZsGreen+ | | 4,500 | 20,000 | 6,998 | 4,856 (69.4) |
| | ZsGreen- | | 3,000 | 10,000 | 4,135 | 3,786 (91.6) |

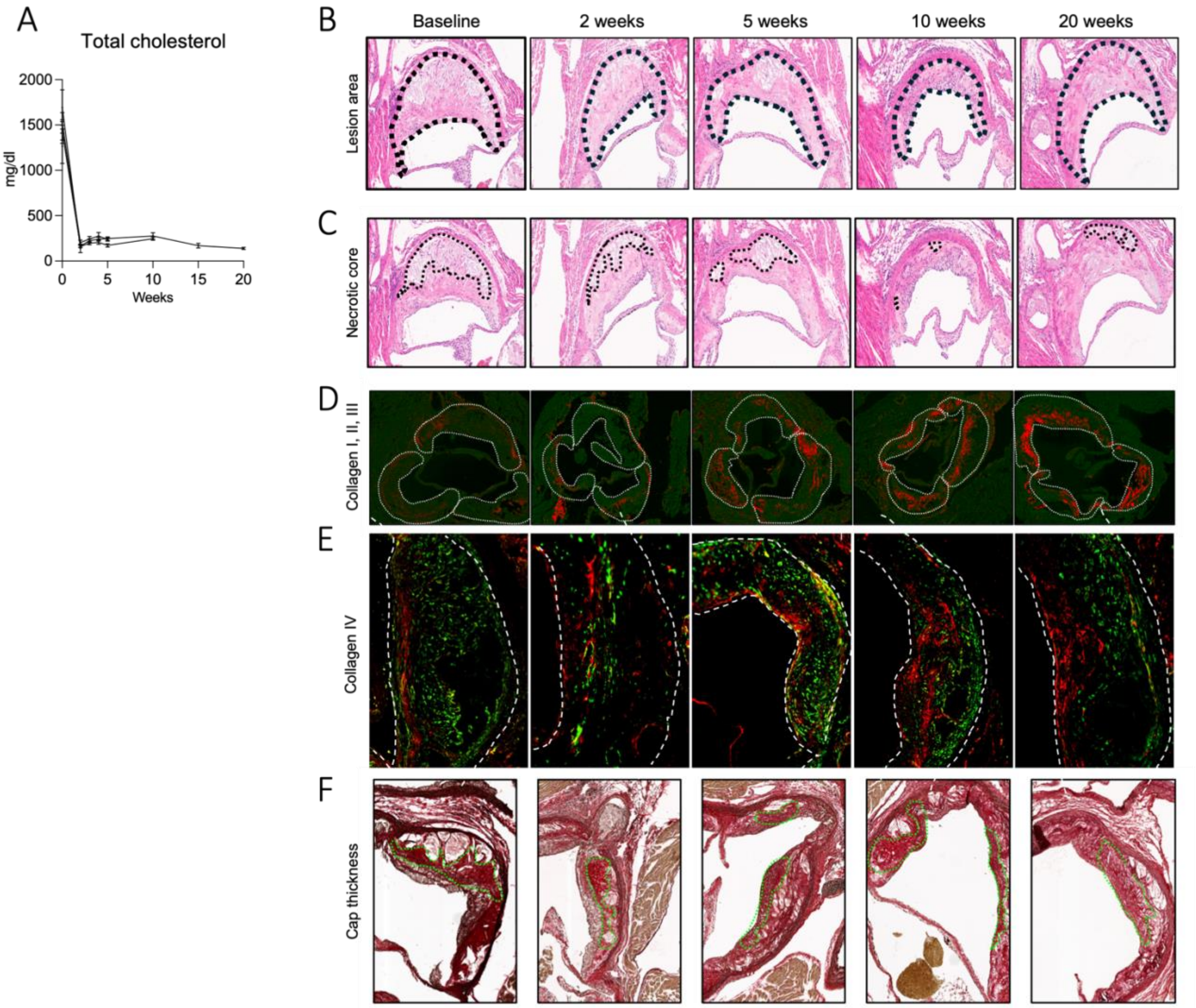


**Supplemental Figure 1. Sustained LDL-C lowering reduces plasma total cholesterol and promotes compositional plaque remodeling. (A)** Plasma total cholesterol concentrations at baseline and throughout 20 weeks of regression. **(B)** Representative hematoxylin and eosin-stained aortic-root sections showing total lesion area. **(C)** Representative hematoxylin and eosin-stained aortic-root sections showing necrotic core area. **(D)** Representative fluorescence images of collagen I, II, and III staining in aortic-root lesions. **(E)** Representative fluorescence images of collagen IV staining in aortic-root lesions. **(F)** Representative Picrosirius Red-stained aortic-root sections illustrating fibrous-cap thickness. Representative images are shown at baseline and after 2, 5, 10, and 20 weeks of regression.

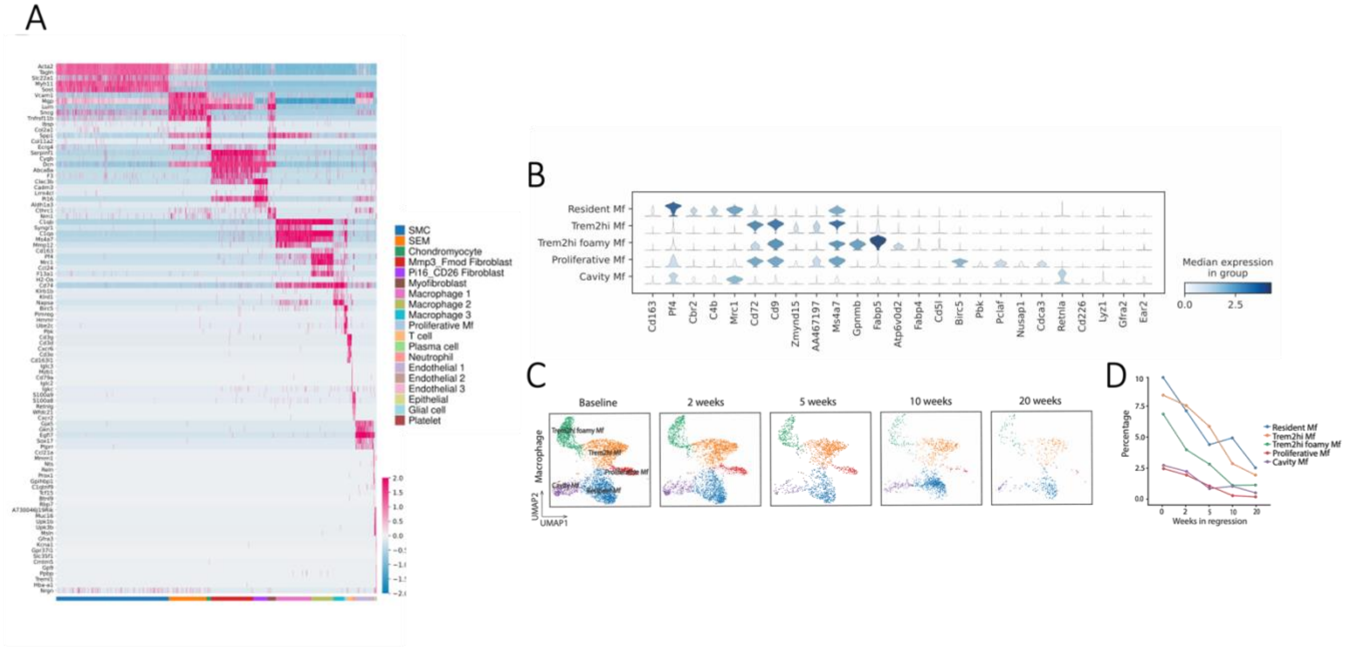


**Supplemental Figure 2. Single-cell profiling reveals distinct cellular populations and macrophage heterogeneity during plaque regression.** **(A)** Heatmap showing scaled expression of representative differentially expressed genes across cell populations identified by scRNA-seq. Columns represent individual cells grouped by population, and rows represent selected marker genes. **(B)** Violin plots showing expression of selected marker genes across five macrophage subpopulations, including resident, Trem2hi, Trem2hi foamy, proliferative, and cavity macrophages. **(C)** UMAP distributions of macrophage subpopulations at baseline and after 2, 5, 10, and 20 weeks of regression. **(D)** Relative proportions of the five macrophage subpopulations across regression time points.
